# How Resource Heterogeneity and Social Threat Shape Intergroup Tolerance: Insights from a Spatial Agent-Based Model

**DOI:** 10.64898/2026.02.21.707157

**Authors:** Cyril C. Grueter

**Author notes:** Corresponding author: Cyril C. Grueter.

## Abstract

The emergence of tolerance between distinct social groups is a central question in social evolution, with relevance to both nonhuman primates and early human societies. Ecological factors such as resource heterogeneity and social threats like bachelor male presence have each been proposed as drivers of intergroup proximity, but their combined effects remain unclear. We developed a spatially explicit agent-based model to examine how resource patchiness and bachelor threat jointly shape aggregation dynamics and intergroup tolerance among mixed-sex groups. In the model, groups forage across heterogeneous landscapes and expand their ranges with increasing patchiness. Bachelor males roam independently and pose localized threats, prompting groups to aggregate probabilistically according to threat intensity. Aggregation decisions follow a sigmoid response, and familiarity accumulates through repeated overlap or joint aggregation. Intergroup tolerance thus arises from encounter histories rather than being preprogrammed. Simulations show that resource heterogeneity promotes tolerance by increasing overlap and encounter frequency, while bachelor threat induces aggregation as a protective response. Tolerance can also emerge without consistent aggregation, provided that ecological conditions repeatedly bring groups together. When both heterogeneity and threat are high, aggregation and familiarity peak, indicating a synergistic effect. These outcomes are robust across a wide parameter space and do not require explicit coordination or cooperative intent. Our findings highlight how simple behavioral rules embedded in ecological and social contexts can yield complex intergroup outcomes, offering a general framework for understanding the evolution of intergroup tolerance in primates and humans.

## Introduction

Understanding the conditions that foster intergroup tolerance is a central challenge in the study of social evolution. While individuals in many primate species live in stable social units, these units are rarely socially or spatially isolated. Instead, groups frequently encounter their neighbors within broader populations or metacommunities. The nature of these encounters varies widely, from aggression and territorial defense to peaceful co-feeding, information exchange, and even affiliative behaviors (Cheney, 1987; Van Belle, Grueter, & Furuichi, 2020). Such variation is increasingly recognized as central to understanding social complexity, and – particularly in humans – the emergence of cumulative culture (Derex & Boyd, 2015; Spikins, French, John-Wood, & Dytham, 2021).

Game-theoretic models such as the Hawk-Dove framework (Smith & Price, 1973) help explain the conditions under which non-aggressive strategies can evolve. In this model, individuals adopt either a Hawk (aggressive) or Dove (non-aggressive) strategy when competing over resources. Doves avoid escalation, thereby minimizing the risk of costly fights. When the costs of aggression are high relative to the value of the contested resource, Dove strategies become evolutionarily stable. Applied to intergroup dynamics, this suggests that peaceful or tolerant interactions may prevail when conflict is too costly or when groups derive mutual benefits from coexistence.

In primates, non-aggressive intergroup interactions can take several forms, ranging from tolerance and brief intermingling to more enduring social associations. Short-term associations have been observed in species such as *Saguinus nigricollis* (Izawa, 1978), *Pan paniscus* (Samuni & Surbeck, 2023), *Gorilla gorilla gorilla* (Forcina et al., 2019), and *Hylobates lar* (Reichard & Sommer, 1997). Longer-term associations, characteristic of multilevel societies, occur in species such as *Rhinopithecus* spp. (Kirkpatrick & Grueter, 2010), *Nasalis larvatus* (Matsuda et al., 2024), *Colobus angolensis ruwenzorii* (Miller et al., 2020; Stead & Teichroeb, 2019), *Papio hamadryas* (Kummer, 1968), and *Theropithecus gelada* (Dunbar & Dunbar, 1975; Snyder-Mackler, Beehner, & Bergman, 2012).

These interactions may be adaptive, with affiliative behavior across group boundaries conferring fitness benefits and being favored by selection under appropriate ecological and social conditions (Furuichi, 2020; Pisor & Surbeck, 2019). Peaceful engagement with outgroup individuals can offer several advantages, including facilitating dispersal (e.g., (Moscovice, Hohmann, Trumble, Fruth, & Jaeggi, 2022)), enabling pre-dispersal prospecting and reconnaissance (e.g., (Sakamaki & Tokuyama, 2024; Sicotte & Macintosh, 2004), reducing predation risk (e.g. (Iwamoto, Mori, Kawai, & Bekele, 1996)), improving access to resources (e.g., (Lucchesi et al., 2020)), promoting reciprocal use of resources (Jaeggi, Boose, White, & Gurven, 2016), and enhancing protection from social threats such as harassment or infanticide by bachelor males (C. C. Grueter & van Schaik, 2010).

Protection from social threats is an important factor influencing grouping behavior, with risks such as male harassment, takeover attempts, and infanticide shaping social dynamics in many primate species (Treves & Chapman, 1996). In many primate species, bachelor males. i.e., mature individuals unaffiliated with a reproductive group, represent a persistent threat to group cohesion and offspring survival. These males may attempt to infiltrate groups, challenge resident males, or commit infanticide. As a counterstrategy, groups may form temporary or more enduring associations with others, enhancing collective vigilance and deterrence capacity (C. C. Grueter & van Schaik, 2010); see also (Rubenstein & Hack, 2004)).

A key ecological factor thought to influence intergroup relations is resource distribution. When resources are evenly spread across the landscape, groups can typically satisfy their needs within small, exclusive territories. In contrast, when resources are patchily distributed, groups may need to expand their ranging area or access shared zones to meet their energetic demands. This increases the likelihood of home range overlap and, consequently, intergroup encounters (Cyril C Grueter, 2022; C. C. Grueter & Pozzi, 2025; C. C. Grueter & White, 2014; Macdonald & Johnson, 2015). Despite its importance, a large-scale comparative analysis directly linking habitat heterogeneity to home range overlap is still lacking. A second key prediction of the Habitat Heterogeneity Hypothesis posits that greater home range overlap should be associated with higher intergroup tolerance. In habitats where exclusive access to key resources is unfeasible, tolerance may be the more economical strategy compared to costly territorial defense. Supporting this idea, a recent cross-species comparative study found that primate species with greater spatial overlap among groups tend to exhibit fewer agonistic intergroup interactions (C. C. Grueter & Lüpold, 2024).

While habitat heterogeneity combined with the threat posed by bachelor males may be sufficient to induce tolerance between groups, aggregation driven by risk avoidance or shaped by landscape features can have important secondary social effects. Prolonged proximity to neighboring groups may lead to the development of intergroup **familiarity**, and with repeated exposure, this familiarity can promote tolerance or even affiliative interactions. When such familiarity, and the resulting reduction in conflict, emerges from sharing a territorial boundary, it is referred to as the “dear enemy” phenomenon (Christensen & Radford, 2018; Getty, 1987).

Despite the intuitive plausibility of these hypotheses, we lack formal models that explore the joint effects of resource heterogeneity and social threat on intergroup tolerance and aggregation. Moreover, few models account for the role of experience-dependent familiarity in mediating these processes. To address these gaps, we developed a spatially explicit agent-based model in which ten mixed-sex groups inhabit a landscape characterized by varying degrees of resource heterogeneity and bachelor male threat. Groups expand their home ranges in response to patchy resources, encounter bachelor males that move randomly across the landscape, and make aggregation decisions based on localized threat. Crucially, intergroup tolerance is not hard-coded: it emerges from repeated spatial overlap, with familiarity increasing over time and influencing subsequent aggregation decisions and co-residence duration. This model allows us to test three key hypotheses: i) That resource heterogeneity promotes tolerance by increasing home range overlap and encounter frequency; ii) That bachelor threat increases group aggregation as a protective response; and iii) That familiarity through repeated encounters amplifies tolerance, even in the absence of direct cooperation.

By integrating ecological and social pressures with a history-sensitive tolerance mechanism, the model provides a formal framework for examining the minimal conditions under which intergroup tolerance can evolve from repeated, risk-induced proximity. It incorporates spatial movement, threat-sensitive aggregation, and familiarity-weighted tolerance, allowing for emergent patterns of social organization shaped by ecological structure and social risk.

## Methods

### Model Overview

We developed a spatially explicit agent-based model in Python 3.10 (Python Software Foundation, 2024) to investigate how resource heterogeneity, bachelor male threat, and familiarity influence the emergence of intergroup tolerance and aggregation among ten mixed-sex groups. The model simulates group interactions over time on a dynamic two-dimensional toroidal grid, where agents forage based on local resource availability, respond to nearby bachelor threats, and update behavioral tendencies based on both current environmental stimuli and accumulated social experience. Simulations were implemented using standard Python libraries, notably NumPy (Harris et al., 2020) for efficient numerical computation and array operations, and Python’s built-in random module for stochasticity in agent initialization and decision-making. This lightweight but flexible computational framework allows tolerance and aggregation to emerge fromsimple, local rules embedded in structured ecological and social contexts, without requiring prespecified affiliative behavior or explicit coordination mechanisms.

### Landscape and Resource Distribution

To simulate variable ecological conditions, we implemented a fixed 50 × 50 toroidal grid. Each cell (*x,y*) represents a spatial patch with a scalar resource value *R*_*xy*_ that influences agents’ foraging behavior. The toroidal topology ensures that agents moving off one edge reappear on the opposite side, eliminating boundary effects.

Resource values were drawn from a beta distribution:

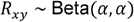

The shape parameter *α* was defined by a heterogeneity parameter *ρ* ∈ [0,1] as:

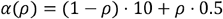

Higher *ρ* values yield patchy resource distributions (low *α*), while lower *ρ* values produce uniform landscapes (high *α*). Resource values were normalized to the interval [0,1] and remained constant throughout each simulation. Groups moved toward more resource-rich cells based on this landscape.

### Agents and Movement

We modeled two types of agents: ten mixed-sex groups and up to five bachelor groups. Each agent occupied a single cell and had discrete spatial coordinates (*x*_*i*_ (*t*), *y*_*i*_ (*t*)) at time *t*.

**Mixed-sex groups** followed a greedy foraging strategy, evaluating all eight neighboring cells (the Moore neighborhood) and moving to the one with the highest resource value:

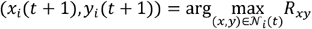

This rule allowed groups to exploit local resource maxima without requiring long-range planning.

**Bachelor groups** moved one step per time unit toward the nearest social group. The toroidal distance was calculated as:

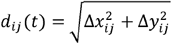

Bachelor groups moved in the direction that minimized *d*_*ij*_ (*t*), simulating male pursuit or social monitoring behavior.

### Aggregation Behavior

At each time step *t*, each social group *i* ∈ {1,…,*N*_*G*_} assesses the risk posed by bachelor groups in its local environment and decides whether to aggregate with other groups. Aggregation is defined as a temporary shift in location toward spatial overlap with other mixed-sex groups in response to perceived threat. Each group defines a threat detection radius *r* that scales with the degree of environmental heterogeneity *ρ*:

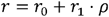

where *r*_0_ =5 is the baseline detection range and *r*_1_ = 10 amplifies sensitivity in patchier environments.

The immediate threat to group *i* at time *t* is quantified as the number of bachelor groups within radius *r*:

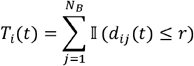

Here, *d*_*ij*_ (*t*) is the toroidal distance between group *i* and bachelor group *j*, and 𝕀 is the indicator function. This threat score is normalized by the total number of bachelor groups to obtain a relative threat level:

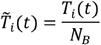

Groups use a sigmoidal decision function to map perceived threat to aggregation probability:

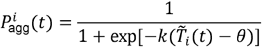

where: *k* controls the steepness of the response curve (default: *k*= 10), and *θ* is the threat threshold for aggregation (default: *θ*= 0.2). This formulation allows for graded, probabilistic responses rather than binary switching.

A uniform random number *u*_*i*_ (*t*) ∼ 𝒰(0,1) is drawn independently for each group and time step. The group aggregates if:

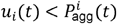

Aggregated groups temporarily shift toward other nearby groups, increasing their likelihood of spatial overlap and repeated co-occurrence. For each group, an aggregation flag *A*_*i*_ (*t*) ∈ {0,1} is recorded at each time step. The aggregation ratio for group *i* is then defined as:

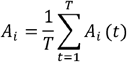

and averaged across groups to yield a global measure of aggregation for the simulation.

### Intergroup Tolerance and Familiarity Matrix

To capture the emergence of intergroup tolerance as a function of repeated spatial proximity, we implemented a familiarity matrix *F*_*ij*_ (*t*), where each entry tracks the familiarity of group *i* with group *j* at time step *t*. Familiarity values evolved dynamically according to spatial co-occurrence. At each time step *t*, for all group pairs *i*… *j*, we updated familiarity using an exponential decay function combined with a spatial overlap condition:

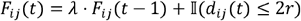

where: *λ* ∈[0,1] is the decay parameter (default: *λ*= 0.99), *d*_*ij*_ (*t*) is the toroidal distance between groups *i* and *j* at time *t, r* is the aggregation radius (as defined earlier), and 𝕀 is the indicator function returning 1 if the two groups are within 2*r*, and 0 otherwise. This formulation ensures that familiarity increases with repeated spatial proximity, and decays when groups no longer encounter each other. The familiarity matrix was assumed to be symmetric:

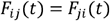

This assumes that familiarity is mutual and accumulates equally for both groups upon co-occurrence.

At the final time step *T*, we computed a group-specific intergroup tolerance score as the sum of familiarity with all other groups:

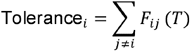

This scalar metric captures the cumulative experience-dependent exposure each group has had to others across the simulation, weighted by both frequency and recency of overlap. To summarize tolerance at the simulation level, we computed the mean intergroup tolerance across all *N*_*G*_ groups:

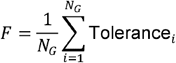

This outcome was used to assess how ecological structure (*ρ*) and bachelor threat influenced the emergent potential for between-group affiliation or peace.

### Emergent Bachelor Threat

Bachelor male threat was modeled as a dynamic, location-dependent social pressure. At each time step *t*, every social group *i* ∈ {1,…,*N*_*G*_} computed its instantaneous threat *T*_*i*_ (*t*), defined as the number of bachelor groups located within its detection radius *r*:

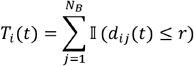

where: *N*_*B*_ is the total number of bachelor groups, *d*_*ij*_ (*t*) is the toroidal distance between social group *i* and bachelor group *j*, and 𝕀 is the indicator function. To summarize threat across time, we computed each group’s mean threat:

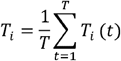

and then averaged over all mixed-sex groups to obtain a simulation-level emergent bachelor threat:

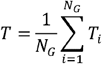

This emergent measure captures both the intensity and spatial persistence of bachelor pressure experienced by groups throughout the simulation, and serves as an input to statistical analyses and visualizations.

### Parameter Space and Replication

We systematically varied two main ecological and social input parameters: Resource heterogeneity: *ρ* ∈ {0.1,0.2,0.3,0.4,0.5,0.6,0.7,0.8,0.9}; Number of bachelor groups: *N*_*B*_ ∈ {0,1,2,3,4,5}. This yielded a total of 5×6 = 30 unique parameter combinations. For each condition, simulations were repeated across multiple random seeds (default: 10 replicates per condition), and all output variables were averaged across replicates.

To assess the robustness of model dynamics, we conducted a one-at-a-time sensitivity analysis, varying key behavioral parameters while holding others constant: Group detection radius: *r*∈ {5,10,15}; Familiarity decay rate: *λ* ∈ {0.95,0.97,0.99,1.0}; Sigmoid steepness: *k*∈ {5,10,15}; Aggregation threshold: *θ* ∈ {0.1,0.2,0.3,0.4}. Outputs were aggregated for each parameter configuration to confirm that observed outcomes were not artifacts of a narrow parameterization.

### Outcome Measures

At the end of each simulation run, we extracted the following primary outcome variables:

#### Aggregation Ratio *A*

For each group *i*, we tracked whether it chose to aggregate at each time step (*A*_*i*_ (*t*) ∈ {0,1}) and computed the average across time:

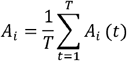

The mean aggregation ratio across all groups was then:

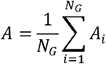

#### Intergroup Tolerance *F*

As detailed above, this metric was derived from the final state of the familiarity matrix and reflects accumulated overlap exposure.

#### Emergent Bachelor Threat *T*

Defined as the simulation-wide average number of bachelor groups encountered per group per time step.

All three outcomes were summarized for each parameter condition. A representative spatial configuration of the model environment, including group home ranges, overlap zones, and bachelor groups, is shown in Figure 1.

**Figure 1.**
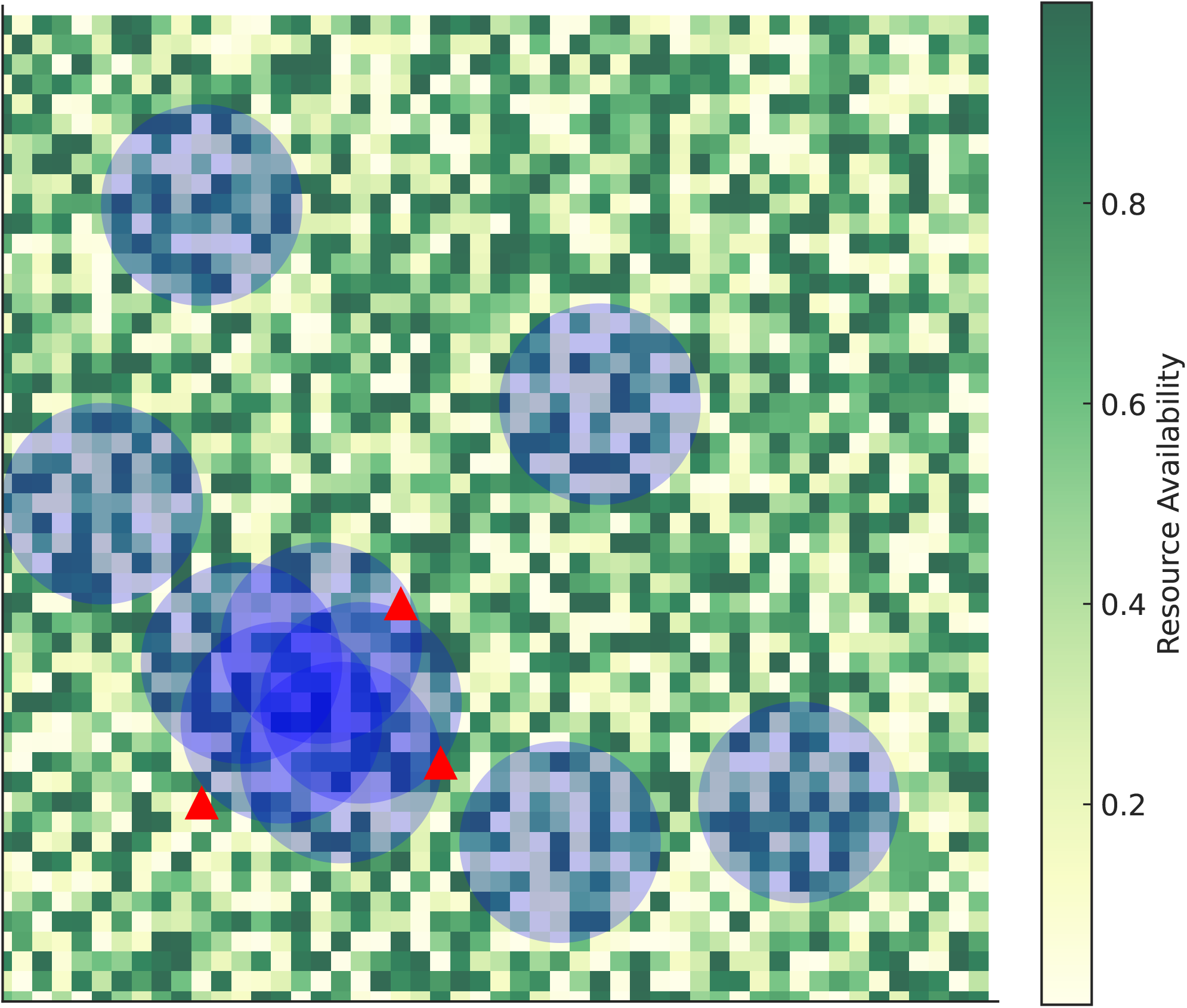
Spatial arrangement of mixed-sex groups, bachelor groups, and resource heterogeneity in the agent-based model. Visualization of ten mixed-sex groups on a 50×50 grid landscape characterized by high resource heterogeneity (ρ = 0.9). Resource availability is shown in light green tones, with brighter areas indicating richer patches. Transparent blue circles represent group home ranges. A subset of five groups in the lower-left quadrant exhibits moderate home range overlap and spatial clustering, forming an aggregated grouping in response to bachelor male threat. Bachelor groups are marked by large red triangles and positioned around the periphery of the aggregated zone, simulating localized social pressure. The figure illustrates how spatial patchiness and external threats jointly shape group proximity and the potential emergence of intergroup tolerance.

## Results

We found that both bachelor threat and ecological heterogeneity had strong and interacting effects on intergroup aggregation and the emergence of tolerance across a broad range of simulated conditions.

### Aggregation Behavior

Aggregation ratios increased systematically with both the number of bachelor groups and the degree of resource heterogeneity. In landscapes with uniform resource distribution and minimal social threat, mixed-sex groups rarely aggregated, maintaining independent foraging trajectories with little spatial overlap. However, as bachelor group presence increased, aggregation became more frequent, especially in ecologically heterogeneous environments.

This effect appears to be driven by two complementary mechanisms: first, an increase in the local presence of bachelor groups raised the perceived threat level within the decision radius of mixed-sex groups; second, patchier landscapes funneled groups toward shared high-value foraging zones. These overlapping pressures increased the probability of both spatial proximity and shared movement, thereby amplifying aggregation frequency. Aggregation was particularly high in simulations where both bachelor threat and resource heterogeneity were elevated, consistent with a synergistic interaction between ecological complexity and social danger.

Heatmaps and scatterplots of model outcomes revealed a clear gradient (Fig. 2a): aggregation ratios remained low at low levels of threat and heterogeneity, increased modestly under moderate conditions, and surged dramatically in the high-threat, high-heterogeneity regime. Trendlines showed that beyond a moderate threshold of threat exposure, the likelihood of aggregation rose steeply, confirming that emergent bachelor pressure plays a central role in triggering group-level defensive convergence.

**Figure 2.**
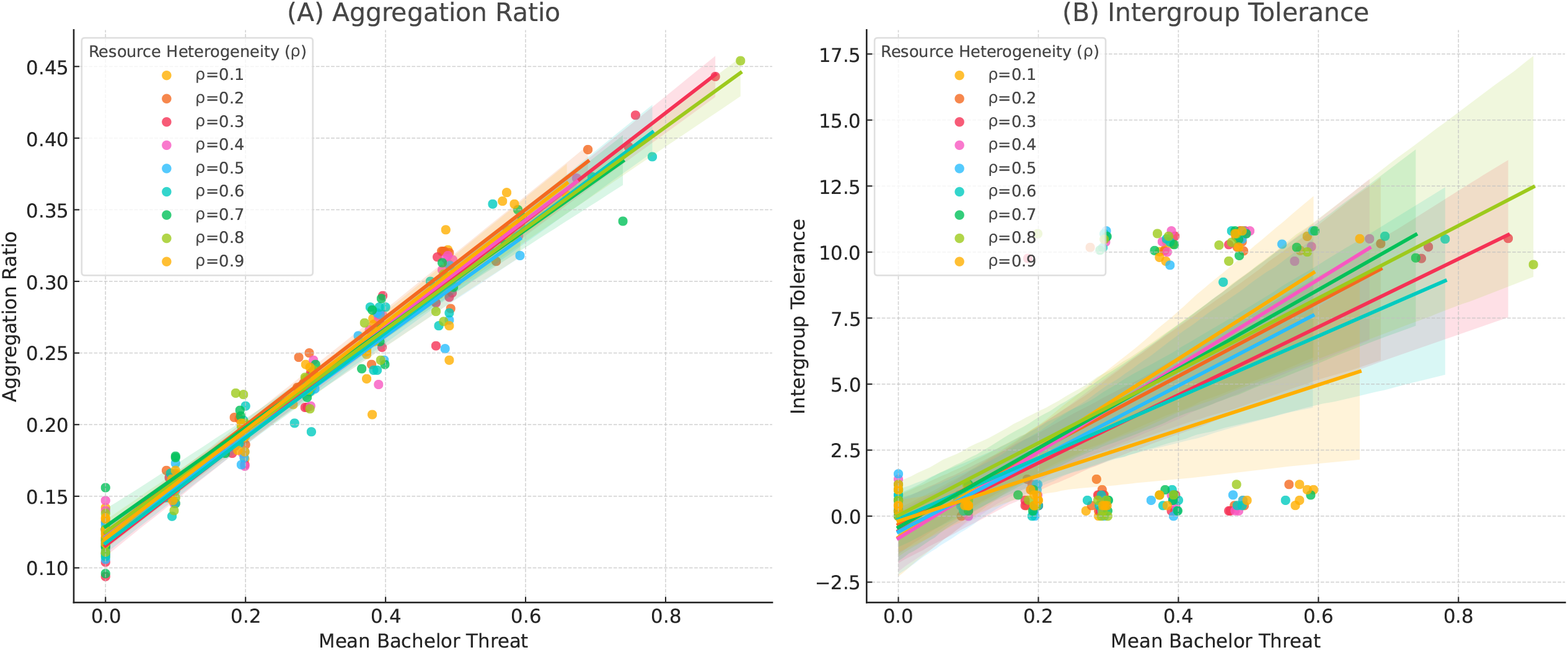
Relationship between mean bachelor threat and (A) aggregation ratio and (B) intergroup tolerance. Each line represents a separate value of resource heterogeneity (ρ), which modulates both social and ecological constraints in the simulation. Aggregation ratio refers to the proportion of time steps during which a group was involved in an aggregation event. Intergroup tolerance is operationalized as mean dyadic familiarity across all mixed-sex groups. The model shows that both ecological heterogeneity and social threat contribute to elevated levels of group aggregation and tolerance.

### Intergroup Tolerance

Patterns of intergroup tolerance, indexed by mean dyadic familiarity, broadly mirrored the dynamics of aggregation (Fig. 2b). However, tolerance was also shaped by the temporal regularity and recurrence of group encounters. In the model, familiarity was allowed to accumulate if groups either shared space or aggregated. This led to more consistent tolerance accrual even when explicit aggregation was intermittent.

Importantly, resource heterogeneity played a critical enabling role: in landscapes with high heterogeneity, groups were more frequently channeled into shared high-quality zones, increasing the chances of repeated spatial proximity. When coupled with bachelor-induced threat, these patchy landscapes created ideal conditions for sustained co-occurrence and familiarity build-up. By contrast, in ecologically homogeneous environments, familiarity scores remained low unless bachelor threat levels were extremely high, suggesting that heterogeneity helps amplify and prolong the spatial convergence needed for tolerance to develop.

Tolerance remained low in simulations with low bachelor threat, even when resource heterogeneity was high. This suggests that ecological complexity alone was insufficient to produce meaningful intergroup familiarity without the additional forcing of social pressure. In contrast, under conditions of frequent bachelor threat, tolerance scores increased markedly. This was driven not by intrinsic social motivation but by the sustained and repeated proximity of mixed-sex groups forced into shared space by external stressors.

These findings support the idea that intergroup tolerance can emerge as a byproduct of defensive behavioral dynamics. When familiarity is allowed to build through either spatial overlap or joint aggregation, social bonds between groups arise as a secondary effect of coordinated threat avoidance, without requiring preexisting affiliative tendencies.

### Sensitivity Analysis

Sensitivity analyses confirmed the robustness of the main results and clarified the conditions under which aggregation and tolerance were most likely to emerge. Aggregation ratio and intergroup tolerance were both most sensitive to changes in the number of bachelor groups and the heterogeneity of the resource landscape. Higher values in either parameter led to consistently higher levels of group convergence and familiarity accrual.

Because familiarity in the model could be incremented by either spatial overlap or co-aggregation, mixed-sex groups in high-threat or high-heterogeneity environments often experienced repeated co-occurrence, even if they did not always aggregate explicitly. Nonetheless, the strongest tolerance scores still arose under conditions of sustained aggregation, reinforcing the importance of prolonged contact in shaping intergroup relationships.

Together, these results highlight a key insight: peaceful intergroup relations can emerge not from intentional cooperation, but from the dynamics of shared risk and overlapping ecological goals. The model provides a plausible pathway for the evolution of intergroup tolerance rooted in defensive convergence, rather than mutualistic intent.

## Discussion

This study used a spatially explicit agent-based model to investigate how ecological and social pressures shape the emergence of intergroup tolerance and aggregation among distinct groups. We found that resource heterogeneity and bachelor male threat independently and interactively shape spatial decision-making and intergroup outcomes: while heterogeneity primarily promoted intergroup tolerance by increasing spatial overlap and familiarity, bachelor threat reliably triggered probabilistic aggregation based on local risk. These findings provide theoretical support for the hypothesis that ecological constraints and social threats act as dual pressures promoting flexible intergroup relationships in primates and other social animals.

Consistent with previous theoretical expectations (e.g., (Cyril C Grueter, 2022; C. C. Grueter & Pozzi, 2025; Macdonald & Johnson, 2015)), our results show that when resources are spatially heterogeneous, groups must expand their home ranges to access necessary resources. This spatial expansion increases overlap with neighboring groups, creating conditions for repeated encounters. By incorporating a familiarity matrix that tracks encounter history, we allowed tolerance to emerge not merely from spatial coincidence, but from repeated spatial overlap or aggregation events (Christensen & Radford, 2018). As a result, groups with high levels of historical overlap were more likely to tolerate one another, even in contexts where current resource competition might suggest conflict.

In contrast, bachelor threat primarily shaped group-level aggregation behavior. Groups reliably aggregated when bachelor males intruded into their home ranges, consistent with the idea that spatial proximity can serve as a form of mutual protection or deterrence (C. C. Grueter & van Schaik, 2010; Rubenstein & Hack, 2004). Interestingly, the combination of high resource heterogeneity and bachelor threat produced synergistic outcomes: not only did groups aggregate more frequently, but they also became more tolerant over time due to an increased number of repeated co-occurrence or aggregation events. These patterns suggest that emergent spatial clustering due to threat, when sustained under ecologically constrained conditions, can facilitate a transition from incidental overlap to structured tolerance.

This study has focused primarily on patterns of intergroup tolerance and association. A crucial next step is to examine the conditions under which such tolerance may transition into active intergroup cooperation. While group aggregation in response to conspecific threats likely represents a protective strategy, i.e., reducing the risk of male takeover and subsequent infanticide through a ‘safety in numbers’ effect, repeated tolerant interactions may lay the groundwork for more coordinated behavior across group boundaries. Such cooperation may manifest in joint action against external threats, in particular conspecific rivals. Evidence for between-group cooperation has been documented in primate species exhibiting multilevel social organization, such as snub-nosed monkeys and geladas. In golden snub-nosed monkeys (*Rhinopithecus roxellana*), for example, multiple resident males from different one-male units have been observed engaging in coordinated patrolling and vigilance, seemingly to deter bachelor males from the surrounding all-male group (Qi et al., 2014; Xiang et al., 2014). Similarly, in geladas, leader males from separate core units have occasionally been reported to jointly repel bachelor males (Mori, 1979; Pappano, Snyder-Mackler, Bergman, & Beehner, 2012; Wrangham, 1976).

Resource heterogeneity may do more than promote tolerance and association; it could also foster intergroup cooperation. (Rodrigues, Barker, & Robinson, 2023) developed a model showing that environments with stable and relatively abundant resources tend to support intergroup tolerance, but not necessarily active cooperation. They argue that the shift toward cooperation is more likely when resource abundance is coupled with spatial or temporal heterogeneity, which creates opportunities or incentives for groups to collaborate.

While our model incorporated familiarity as a factor influencing intergroup tolerance, another key determinant may be genetic relatedness. Reduced genetic differentiation between groups, which can arise through mechanisms such as non-random dispersal or extra-group mating, can contribute to elevated tolerance levels (for a review, see (C. C. Grueter & Pozzi, 2025)). Familiarity itself may develop through repeated intergroup encounters, but it can also stem from shared social history, such as prior group membership before events like dispersal or fission (Furuichi, 2020; Mirville et al., 2018; Scarry & Tujague, 2012). Although familiarity and relatedness can independently influence tolerance, they often interact and may be difficult to disentangle in empirical studies. In many cases, familiarity may serve as a proxy for relatedness, or vice versa, with both factors jointly shaping patterns of intergroup behavior.

While our model incorporates several biologically grounded processes, including spatially explicit movement, threat agents, and memory-dependent social behavior, it remains intentionally parsimonious. We abstracted from individual-level decision-making, food depletion, and demographic processes such as birth, death, and dispersal. This was a deliberate choice, grounded in the goal of identifying general mechanisms underlying the emergence of intergroup tolerance and the conditions that lead to intergroup coexistence.

This simplified model aligns with the function of agent-based models as generative tools for theory development (Epstein, 2012). By minimizing assumptions and avoiding species-specific parameterization, it remains broadly applicable across taxa, including non-human primates, early human societies, and other social mammals. Nonetheless, future iterations of the model could incorporate greater ecological and social realism, for example by introducing agents with variable personalities or decision-making styles. Additional factors that may influence whether groups reduce aggression, tolerate one another, or form associations include the relative competitive capacity or resource-holding potential of opposing groups (e.g., (Mirville et al., 2018)), decision-making dynamics within groups (e.g., democratic vs. despotic leadership; (Hunt et al., 2024)), population density (e.g., (Decellieres, Zuberbühler, & León, 2021)), the presence of estrous females (Kitchen, Cheney, & Seyfarth, 2004), and the context or location of encounters (e.g., (Morrison et al., 2020)). Some of these variables could be incorporated into more complex models to refine predictions. In parallel, empirical research can inform and validate key parameters such as how resource patchiness affects home range size, or how often bachelor males intrude in natural populations, thereby improving the model’s relevance and accuracy.

Our findings have several implications. First, they provide formal support for the idea that resource-driven ranging behavior and social threat avoidance can independently and jointly promote intergroup tolerance, especially when reinforced by familiarity through repeated contact. Second, the model suggests that intergroup tolerance may not require cooperative intent at the outset but can emerge through spatial and social constraints. Finally, the results raise interesting questions about how early human groups or ancestral primates may have transitioned from xenophobic to more tolerant intergroup dynamics in response to environmental and social pressures.

## Acknowledgments

I am grateful to Tyler Bonnell for his valuable feedback on this manuscript.

